# Integrating deep and radiomics features in cancer bioimaging

**DOI:** 10.1101/568170

**Authors:** A. Bizzego, N. Bussola, D. Salvalai, M. Chierici, V. Maggio, G. Jurman, C. Furlanello

## Abstract

Almost every clinical specialty will use artificial intelligence in the future. The first area of practical impact is expected to be the rapid and accurate interpretation of image streams such as radiology scans, histo-pathology slides, ophthalmic imaging, and any other bioimaging diagnostic systems, enriched by clinical phenotypes used as outcome labels or additional descriptors. In this study, we introduce a machine learning framework for automatic image interpretation that combines the current pattern recognition approach (“radiomics”) with Deep Learning (DL). As a first application in cancer bioimaging, we apply the framework for prognosis of locoregional recurrence in head and neck squamous cell carcinoma (N=298) from Computed Tomography (CT) and Positron Emission Tomography (PET) imaging. The DL architecture is composed of two parallel cascades of Convolutional Neural Network (CNN) layers merging in a softmax classification layer. The network is first pretrained on head and neck tumor stage diagnosis, then finetuned on the prognostic task by internal transfer learning. In parallel, radiomics features (*e.g.*, shape of the tumor mass, texture and pixels intensity statistics) are derived by predefined feature extractors on the CT/PET pairs. We compare and mix deep learning and radiomics features into a unifying classification pipeline (RADLER), where model selection and evaluation are based on a data analysis plan developed in the MAQC initiative for reproducible biomarkers. On the multimodal CT/PET cancer dataset, the mixed deep learning/radiomics approach is more accurate than using only one feature type, or image mode. Further, RADLER significantly improves over published results on the same data.

## I. Introduction

Artificial Intelligence (AI) progress in medical image interpretation is rapidly gaining speed, with a wide range of applications [1]–[3]. Its translation to clinical practice is expected to accelerate due to faster regulatory approval procedures for medical algorithms [4]. As deep learning models (DL) aim to evolve status from exploratory to clinically effective solutions, interpretability remains a major stepping hindrance [4], [5]. In general terms, DL provides a class of machine learning methods that can model complex abstractions of patterns through multiple non-linear transformations estimated by data-driven training procedures. Convolutional Neural Networks (CNNs) are DL models widely successful in image recognition and classification. Their application in medical image analysis dates back to 1996, to discriminate tumor mass and normal tissue in mammography [6]. Since then, CNNs have provided results comparable to experts in the diagnosis of skin lesions [7], classification of colon polyps [8], [9], survival analysis of glioma [10], ophthalmology [11], histology [12], and other areas [1].

Medical imaging is indeed a key resource shaping the clinical trajectory of a patient. Based on these initial success stories, DL techniques are expected to represent a major breakthrough in diagnosis, treatment decision, prognosis and treatment evaluation. This breakthrough is expected to be pervasive and valid over the diverse medical imaging modalities, *i.e.*, anatomical (such as CT scan) or functional (*e.g.*, PET).

In apparent competition with DL, radiology is already walking fast on a critical innovation path enlightened by *radiomics*, the umbrella-term for pattern recognition methods composed by quantitative image feature extraction paired with statistical or standard machine learning classifiers. Radiomics is grounded on the underlying biological assumption that imaging features can capture distinct phenotype morphology [2], thus achieving both classification and clinical understanding in the machine learning process.

This emphasis on interpretability is a key factor in oncology, where molecular expressions of cancer subtypes may manifest as tissue architecture and nuclear morphological alterations [13]; hence automatic evaluation of disease aggressiveness and patient subtyping can be derived to inform the therapeutic decision. Radiomics features include descriptors of intensity distribution, spatial relationships, texture heterogeneity patterns, descriptors of shape and morphology, and volumetric quantification [14], [15]. Radiomics features can be extracted by tools such as the cancer imaging phenomics toolkit (*CaPTk*) for radiographic images [16], *histomicsTK* for histological Whole Slide Images, or *pyRadiomics* [17].

The role of features in DL is remarkably different: by construction, data are non-linearly mapped throughout the transformation connecting the input and output spaces of a neural network. At each layer, data are projected in a synthetic feature space defined by the training process; such latent features can be investigated in association to the outcome labels. Although hard to define in biological or morphological terms, these learned features can outperform the hand-crafted ones [18]–[21].

However, DL models typically need a much larger amount of data for training for optimal results than statistical machine learning models; thus these models are often bootstrapped With the transfer learning approach, *i.e.* borrowing weights of models trained on different domains, and possibly retraining only a sector of interest of the network with the data from the novel task [22]. This trick is extensively used in non-medical domains, based on the availability of large-scale data and pretrained architectures [23], [24]. Recently, these resources are becoming available in cancer research. For example, the *DeepLesion* dataset, containing over 32, 000 annotated lesions in CT scans [25], and *The Cancer Imaging Archive* (TCIA), which provides medical images of different modalities (MRI, CT, etc.) [26].

The success of transfer learning schemas is clearly contributing to approaching DL models as powerful extractors of useful feature sets (i.e. *deep features*). However, linking deep features to meaningful clinical properties interpretable by physicians remains a key challenge [3]. Statistical machine learning approaches are also still widely used in radiomics [27], [28]. This state of the art has naturally led to the idea of a hybrid combination of hand-crafted radiomics (HCR) and deep-learning radiomics (DLR) in an integrated system system [4], [29], [30]. These systems can provide objective characterizations of tumor and a more effective decision support environment, activating expertise in interpretation by clinicians, biologists and bioinformaticians [31].

Notably, the fusion between the two radiomics feature types operates either at decision level or at feature level. With the first approach, models built on HCR and DLR features are developed separately and a final decision module combines their outputs [19], [32]. With the second approach, the integration of HCR and DLR features operates at early level in a multimodal learning framework (e.g. by concatenation), usually with better classification performance [23], [33]–[37].

In this work, we propose RADLER, an automatic pipeline for the integration of DLR and HCR features for medical images analysis, in a first application on multimodal PET/CT scans. To support reproducibility, models are trained with a Data Analysis Plan (DAP) that includes repeate cdross-validation, model selection and feature ranking techniques. To validate the framework, an application is shown on a dataset of two-modality 3D CT/PET scans for prognosis of locoregional recurrence (LR) in head and neck squamous cell carcinoma (N=298), previously solved with a HCR approach and a logistic regression model [38]. The multimodal network architecture is derived from a multistream multiscale architecture for lung cancer screening [39]. The network is first pretrained on head and neck tumor stage (T-stage) diagnosis, then finetuned on the prognostic task (internal transfer learning). The RADLER model integrates in this case up to four feature types (CT-HCR, CT-DLR, PET-HCR, PET-DLR) improving over the published results on the same data [38]. Moreover the mixed deep learning/radiomics approach is more accurate than using only one feature type, or image mode.

## II. Materials and Methods

### Head-Neck-PET-CT Dataset

The Head-Neck-PET-CT (HN) dataset ^1^ has been originally introduced in [38], and further used in [40]. It includes medical images and clinical data of 298 patients with head and neck squamous cell carcinoma.

For each patient, the HN dataset provides CT and PET scans and Gross Tumor Volume (GTV) mask, preprocessed according to the pipeline in Fig. 1. Several clinical variables are included, in particular the Locoregional Recurrence (LR) within the follow-up period (median: 43 months; range: 6-112 months), and T-stage at diagnosis. Data are gathered from four different hospitals, each one representing a single cohort. Notably, each hospital has its own image acquisition equipment and acquisition settings, which is a cause of heterogeneity in image characteristics, in particular resolution of the PET images. Moreover, The HN dataset is highly unbalanced for the LR prognosis, with 15.8% of recurrence (Table I).

**TABLE I.** HN DATA: CLASS DISTRIBUTION OF THE LR PROGNOSTIC TASK IN TRAIN AND TEST SETS (N=294).

|  | no-LR | LR | total |
| --- | --- | --- | --- |
| Train | 164 | 27 | 191 |
| Test | 88 | 15 | 103 |

**Fig. 1.**
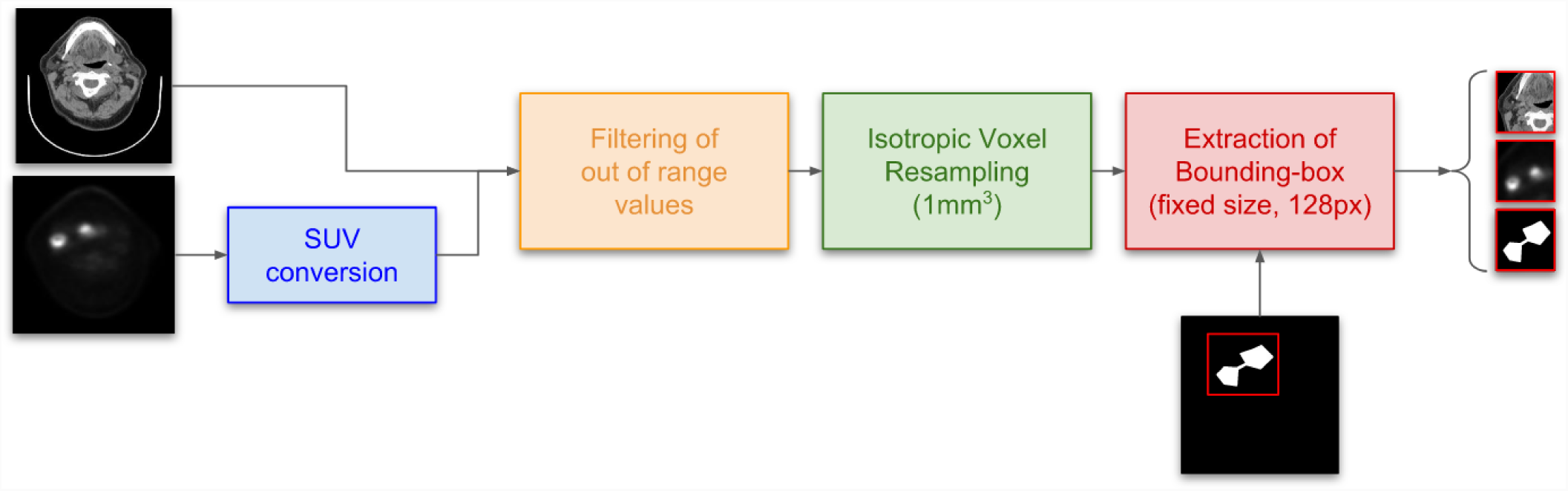
Workflow of CT and PET volumes preprocessing pipeline. SUV: Standardized Uptake Values

For the sake of comparison, we split the HN dataset into training and test sets with the same partition as in the original study [38]. In particular, two hospital cohorts are used to train the model (*training set, N*_*tr*_ = 191), and two cohorts are used for testing (*test set, N*_*te*_ = 103), with proportioned class stratification (see Table I); the design is chosen to consider possible batch effects in the subcohorts due to the hospital of provenance.

The data set includes the secondary diagnostic label tumour stage (T-stage: score in a 1-4 scale), which was considered for the internal transfer learning strategy. Patients missing the T-stage were not considered in training the diagnostic model, thus developed on a subset of 269 patients, partitioned into 60/40% train/test sets (see Table II).

**TABLE II.** HN DATA: CLASS DISTRIBUTION OF THE DIAGNOSTIC T-STAGE TASK IN TRAIN AND TEST SETS (N=269).

|  | T1 | T2 | T3 | T4 | total |
| --- | --- | --- | --- | --- | --- |
| Train | 29 | 64 | 36 | 34 | 163 |
| Test | 10 | 45 | 28 | 23 | 106 |

### Image Processing Workflow

As summarized in the diagram in Fig. 1, CT and PET images are first preprocessed to obtain a standardized input information. In particular, PET images are converted to Standardized Uptake Values (SUVs), applying the protocol proposed by the Quantitative Imaging Biomarkers Alliance (QIBA), which also considers vendor-dependent parameters. It is worth mentioning that the conversion of PET images to SUV format is still an open question [41], [42]. The GTV pattern is imported by creating a binary mask with the same size of the CT and PET images (“GTV mask”). For both CT and PET modalities, the preprocessing pipeline includes: thresholding on the pixel values; isotropic voxel resampling; and extraction of the 3D volume containing the GTV.

The intensity in CT images is associated to tissue density and it is measured with the Hounsfield scale (HS). Specific HS value ranges are defined for each type of tissue, which allows the direct comparison of images from different vendors. However, artifacts or acquisition errors might affect the image and give pixel values outside the physiological range. We filter these artifacts by thresholding the pixel values of CT images between HS=-1050 (air density score) and HS=3050 (bone density score). Similarly, the SUVs in PET images are thresholded in the range between 0 and 50 to avoid artifacts due to errors in sensors readings.

Further, isotropic voxel resampling was performed on CT and PET images, as well as on the GTV mask, based on cubic interpolation of each image on a 3D grid with 1 *mm*^3^ voxels, in order to have an homogeneous standard spatial information.

The last module in the image preprocessing pipeline extracts a subvolume of the image which contains the GTV. This reduction enables to compute the radiomics features only from the voxels, also reducing the size of the 3D image portion to analyze with DL on the Graphical Processing Unit (GPU) memory. The drawback of this operation is the loss of contextual information near the GTW, thus the normalized size of the subvolume was set to 128 *mm*^3^, a reasonable trade-off between the size of the GTVs in the dataset and the amount of context included. The volume of interest was centered in the center of mass of the GTV, also used to center the subvolumes of the CT, PET and GTV mask images.

In summary, the output of the image processing workflow (see Fig. 1) is composed by three 128 *mm*^3^ images, one for each modality and for the GTV mask.

## III. The Radler Integrative Radiomics DAP

We have developed the RADLER radiomics pipeline as a general framework for predictive models that can integrate Deep Learning and predefined features. The framework is also designed to manage multimodal imaging datasets, as in the case of study of LR prognosis on the HN cancer dataset. The main steps in the RADLER pipeline after the preprocessing phase are described below as exemplified on the LR HN task (see Fig. 3.

### Radiomics Feature Extraction and Integration

Three sets of radiomics features are considered:

i. *HCR.* A total of 3, 249 radiomics features are extracted for each patient, replicating [38]. The feature extraction is based on the `pyradiomics` framework [29]. The HCR features are chosen to describe three main image properties: shape (13 features, based on the GTV contours), intensity (18 features, based on the voxel intensities), and texture (1, 600 features, based on four Gray-Level Matrices). Following [38], 40 types of texture features were considered, each one computed on 40 sets of parameters that define pixel spacing, quantization method and number of gray levels;
ii. *DLR.* A total number of 512 deep features are extracted (256 from PET images and 256 from CT images) as a byproduct of a multimodal neural network. The network is trained on CT and PET simultaneously [39], with two identical and parallel convolutional branches merged in a fully connected layer (see Fig. III for the details on the architecture). An internal transfer learning procedure is applied by first training the whole network on the T-stage dataset, then predicting LR by finetuning, *i.e.* retraining only the linear blocks (final blue box in Fig. III). Fixed hyper-parameters are used to regulate the training process with Adam [43] optimizer (batch size: 32, epochs: 500, learning rate 10^-3^). Data augmentation procedures were used to improve the performance and reduce overfitting: i.e., minimal rotations, translations and Gaussian noise. The transformed images were resized to cubes of 64× 64 × 64 to better fit the GPU memory size.
iii. *HCR* + *DLR.* The two types of features are concatenated into an integrative dataset. A more accurate model is expected from two types that should capture different and complementary information from the input images.

### Feature selection and Ranking

The feature selection section in RADLER leverages a combination of three methods from `scikit-learn` [44]. Features are standardized after imputation of missing values (*Nan* and *inf*) by mean feature values. The procedure is composed of three main steps:

- Removal of correlated features (UNCORR). Since the same types of radiomics features are extracted several times with different sets of parameters (e.g. voxel size for interpolation), the HCR feature set includes highly correlated features. Thus, the Pearson’s correlation matrix is computed, and high correlated features (*ρ* > 0.95) are removed;
- Univariate analysis (UA). An association score (ANOVA F-test) is computed between each feature and the target. Features are ranked based on the association score, keeping the top 1, 000 features;
- The remaining features are ranked based on their predictive power within a Recursive Feature Elimination (RFE) procedure and ordered by decreasing importance.

Feature selection and ranking are performed for each feature set type. Notably, no feature from the deep feature sets is removed by the UNCORR step; thus the features automatically created by the network are highly uncorrelated, *i.e.* the information content is maximized. Table III summarizes the three feature sets and the results of the feature selection section.

**TABLE III.** SUMMARY OF THE THREE FEATURE SETS

| Feature Set | # samples | Number of features |  |  |
| --- | --- | --- | --- | --- |
|  |  | Initial | After UNCORR | After UA |
| HCR | 295 | 3249 | 968 | 968 |
| DLR | 294 | 512 | 512 | 512 |
| HCR + DLR | 293 | 1480 | 1427 | 1000 |

### Classification within a Data Analysis Protocol framework

A linear Support Vector Machine (LSVM) model is trained on the three feature sets within a Data Analysis Protocol (DAP) framework. The DAP was derived from a bioinformatic machine learning procedure developed by the MAQC consortium to grant reproducibility of predictive biomarkers from microarrays and next-generation sequencing platforms, thus in a massive data context [45]–[47]. The dataset is split beforehand into training and test sets (two cohorts for each split, see Table I): the training set is used to develop the model and the test set is used only to asses the predictive performance. The Matthews Correlation Coefficient (MCC) [48]–[50] is used as the evaluation metric.

A grid of parameters is created from the values of the LSVM regularization parameter *C* ∈ {10^-3^, 10^-2^, 10^-1^, 1, 10, 100} and the increasing number of features *nf* ∈ { 0.1%, 0.2%, 0.5%, 1%, 2%, 5%, 10%, 20%, 50%, 100%}. For each parameter point, the training set is randomly split into 5 folds, which are cyclically used to train and validate the model. The optimal parameters are selected by maximizing the predictive results on the validation set. This procedure is repeated 10 times, thus obtaining 50 predictive scores for each parameter point, which are averaged and used to select the best parameter set. Finally, the optimal predictive model is trained on the whole training set using the best parameters and evaluated on the test set.

## IV. Results

In order to obtain the Deep Learning network for the LR task, the architecture was first trained to classify the T-stage, with MCC = 0.863 on the training set (one-shot) and MCC = 0.279 on the test set. As this network is only used to initialize the parameters to predict the LR, this result was not validated within the DAP procedure. To qualitatively investigate the embedding resulting in the T-stage network, we considered the Uniform Manifold Approximation and Projection (UMAP) dimensionality reduction method [51]. The UMAP projection for the T-stage data of the Deep Features extracted from the PET images is displayed in Fig. 2 (left); the deep learning model transforms the input images into a representation of the T-stage severity, which can be qualitatively represented as a trajectory in the projection plan (see Fig. 2, right).

**Fig. 2.**
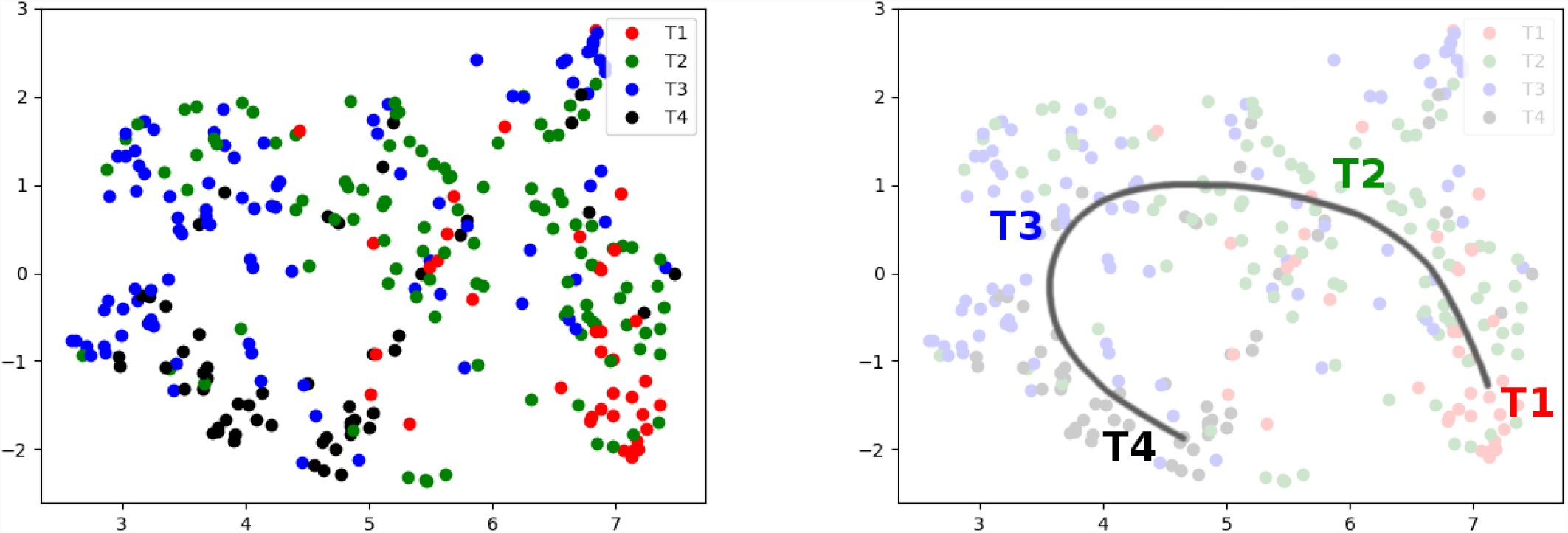
UMAP embedding of the radiomics deep features extracted from the PET images (PET-DLR). Each point represents a patient, colour coded for T-stage. On the right panel, a qualitative trajectory of cancer severity is overlaid on patient clusters of increasing T-stage.

**Fig. 3.**
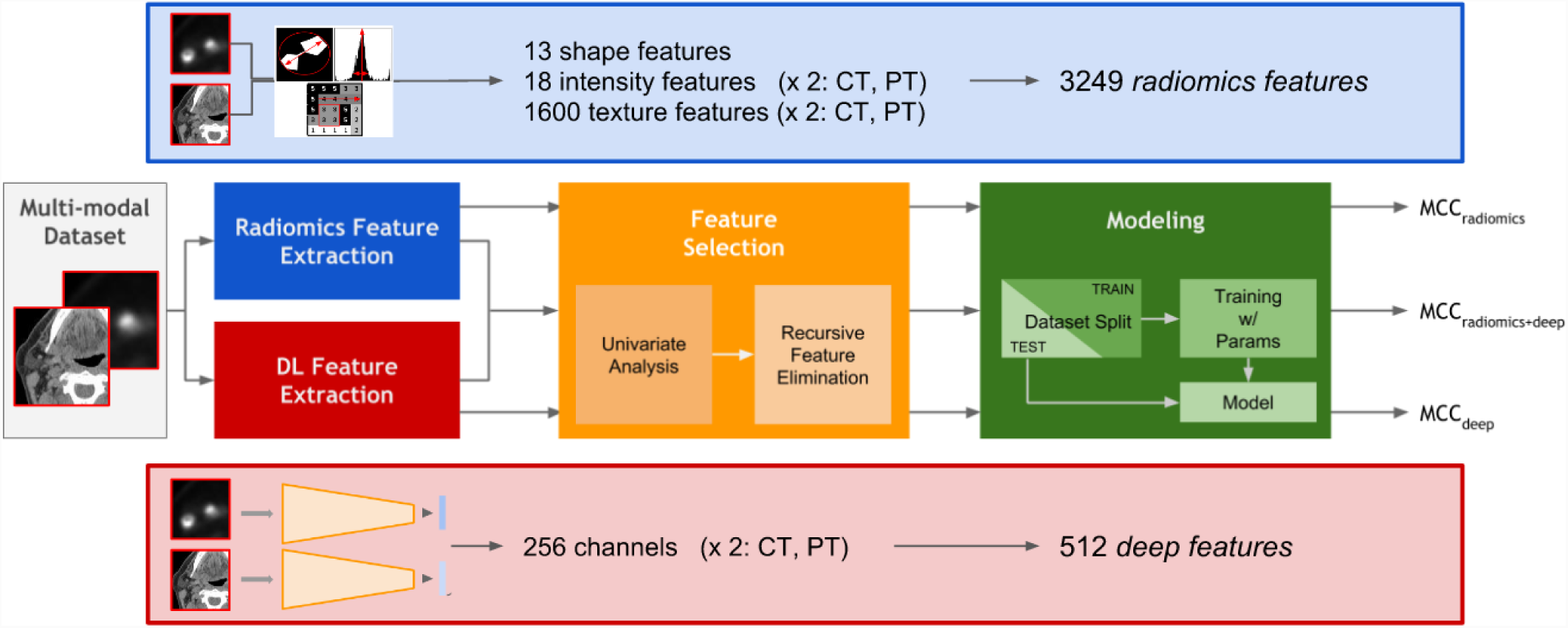
The RADLER pipeline on CT/PET data: predictive models from the integration of radiomics and deep features.

**Fig. 4.**
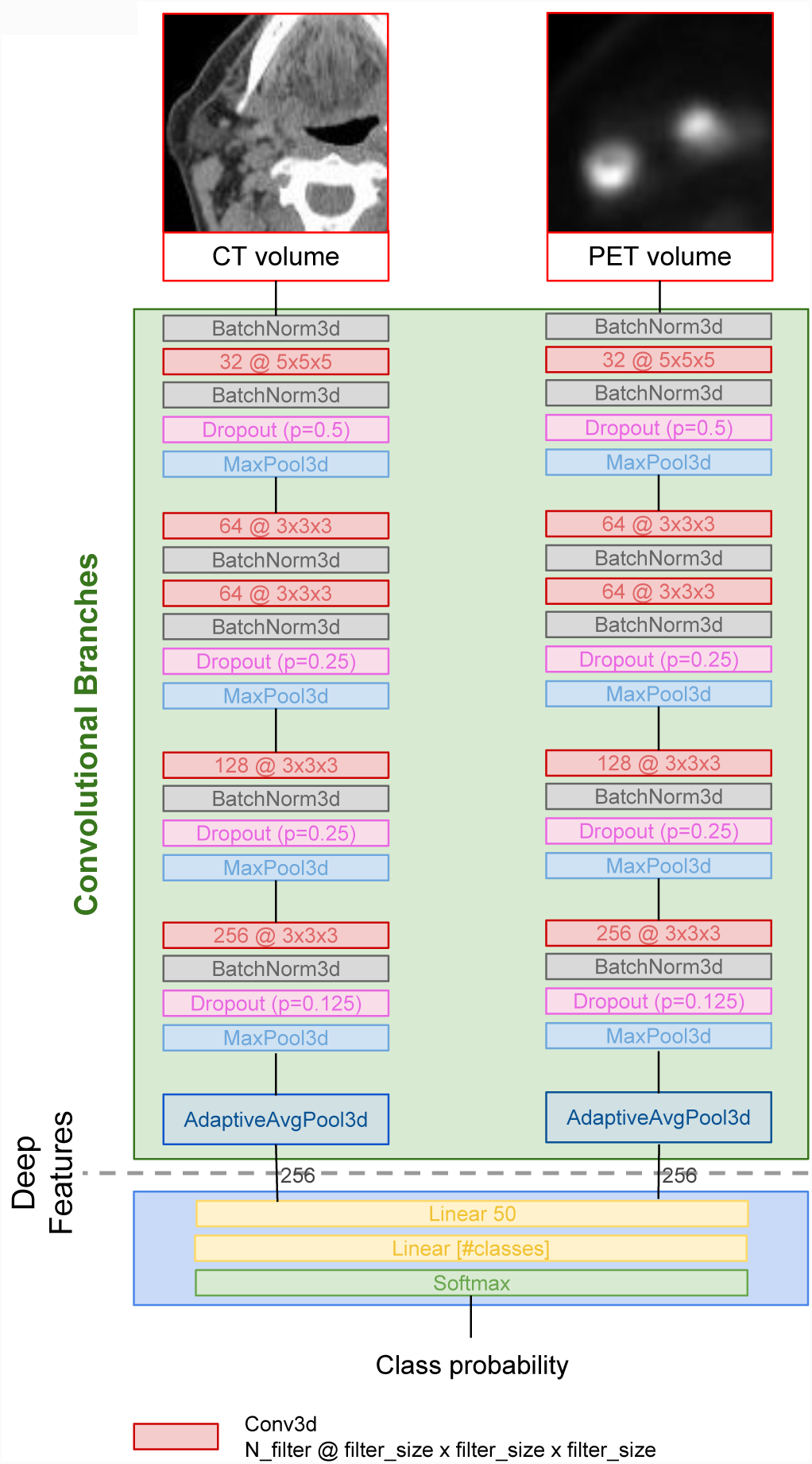
Multimodal network architecture for CT/PET scans. The network inputs are pairs of volumes of size 64 × 64 × 64, one for each channel (CT and PET). The total number of output features from the convolutional branches is 512. The final output of the network is the probability of LR occurrence.

We transferred the weights of the convolutional branches to a new network and trained the linear layers in the final block to predict LR. These branches are used to generate the deep features (DLR feature set). The trained network obtains MCC_train_ = 0.367 and MCC_test_ = 0.245, with a small overfitting.

We compared the performance of the LSVM model trained within the DAP on the different feature sets (see Table IV) and against the reference study [38] (see Table V).

**TABLE IV.**
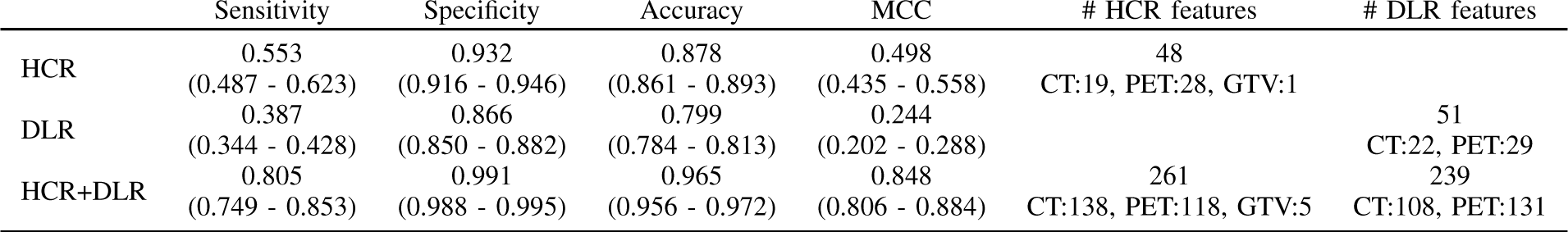
CROSS-VALIDATION PERFORMANCE FOR THE DIFFERENT FEATURE SETS ON THE TRAINING SET, USING THE DAP PROCEDURE. METRICS ARE EXPRESSED AS MEDIAN VALUES WITH 95% BOOTSTRAPPED CONFIDENCE INTERVALS. THE LAST TWO COLUMNS REPORT THE NUMBER OF FEATURES PER CLASS AND MODALITY.

|  | Sensitivity | Specificity | Accuracy | MCC | # HCR features | # DLR features |
| --- | --- | --- | --- | --- | --- | --- |
| HCR | 0.553<br>(0.487 - 0.623) | 0.932<br>(0.916 - 0.946) | 0.878<br>(0.861 - 0.893) | 0.498<br>(0.435 - 0.558) | 48<br>CT:19, PET:28, GTV:1 |  |
| DLR | 0.387<br>(0.344 - 0.428) | 0.866<br>(0.850 - 0.882) | 0.799<br>(0.784 - 0.813) | 0.244<br>(0.202 - 0.288) |  | 51<br>CT:22, PET:29 |
| HCR+DLR | 0.805<br>(0.749 - 0.853) | 0.991<br>(0.988 - 0.995) | 0.965<br>(0.956 - 0.972) | 0.848<br>(0.806 - 0.884) | 261<br>CT:138, PET:118, GTV:5 | 239<br>CT:108, PET:131 |

**TABLE V.** PERFORMANCE FOR THE DIFFERENT FEATURE SETS ON THE TEST SET, COMPARED TO REFERENCE RESULTS (“VALLIÈRES CT-PET”).

|  | Sensitivity | Specificity | Accuracy | MCC |
| --- | --- | --- | --- | --- |
| Vallières CT-PET | 0.56 | 0.67 | 0.65 | Not reported |
| HCR | 0.94 | 0.95 | 0.95 | 0.832 |
| DLR | 0.53 | 0.97 | 0.9 | 0.57 |
| HCR+DLR | 0.67 | 0.91 | 0.94 | 0.748 |

Notably, on the test dataset we improve the original results for all feature sets and metrics, except for the sensitivity achieved using the deep features (0.53 vs 0.56: Table V). In particular, the best results on the test dataset are achieved with the radiomics features (Sensitivity: 0.94; Specificity: 0.95; Accuracy: 0.95).

However, an underfitting effect is observed for both the HCR and DLR feature sets, with higher accuracy on the test than on the training set. The overall best performance is achieved with the HCR+DLR feature set, i.e. MCC_DAP_ = 0.848.

Considering the DLR set, we note that both the linear layers of the DL network and the LSVM model have the same sets of features as input, and so their performance can be compared. Notably, the performance of the linear layers (MCC = 0.245) on the test set are within the confidence intervals of the LSVM model (MCC = 0.244).

We also investigated whether considering both CT and PET is effectively useful, *i.e.* if the two modalities contribute with complementary information (see Table VI). Please note that the single modality feature sets (CT and PET) are obtained from the corresponding multimodality features sets after the feature selection and ranking step, by selecting the features computed from the respective images.

**TABLE VI.**
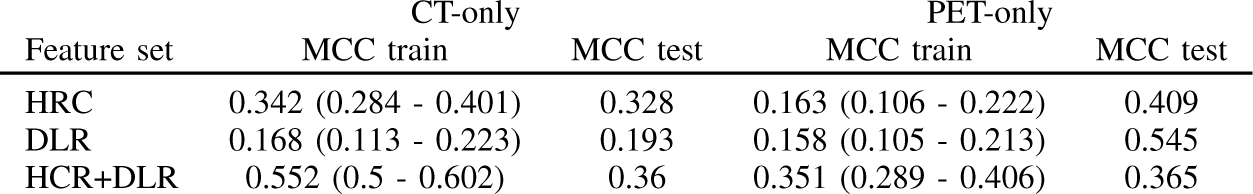
PERFORMANCES ON SINGLE MODALITY FEATURE SETS.

Results reported in Table VI show that the underfitting issues, *i.e.* MCC_test_ ≫ MCC_train_, mostly affect the PET-only modality, thus confirming the intrinsic difficulties of quantitative interpretation of PET images. This is also an open problem also in the clinical practice, in particular for head and neck pathologies [52], [53]. In fact, despite the applied conversion to SUV, technical differences between PET scanners, and non-linear effects associated to GTV segmentation as well as other patient-dependent parameters are not taken into account [41], [42], [54]. Further, we observe that the best MCC is achieved by the HCR+DLR feature set, also resolving the mentioned underfitting issues.

## V. Discussion

The RADLER framework introduced in this study aims at the integration of deep and radiomics features for medical image analysis and classification. Its first application in a prognostic task of locoregional recurrence (LR task) of head and neck cancer improves with respect to the state of art [38], both in terms of sensitivity (0.94 vs 0.56) and specificity (0.95 vs 0.67). Moreover, the DAP included in the framework is used to evaluate variability due to resampling and control for selection bias in the model selection phase. As assessed by the DAP, the feature set integrating radiomics and deep features is more effective in predicting LR than only one of the feature types.

The RADLER framework is demonstrated with a DL architecture for CT/PET; in detail, a 3D multimodal CNN is adapted from a 2D solution originally aimed at classifying lung nodules from CT imaging [39]. Secondly, we adopted an internal transfer learning approach, starting from the diagnostic classification of tumour stage. This domain adaptation approach proved useful in dealing with class unbalance and a relatively low number of samples, while achieving good predictive performance, as shown on the HN dataset, with high class unbalance and low number of samples.

This design makes the RADLER pipeline and its DL network potentially effective to model other clinical tasks in which different image modalities (*e.g.*, MRI) and anatomical regions (*e.g.*, lung, brain) are considered. The pipeline still requires the manual annotation of the GTV; however, this task could be tackled by automatic segmentation models, thus moving towards a fully automated pipeline. The development of a multimodal network in the RADLER framework was driven as a first step by the integration of PET and CT images in a clinical context. Further work is needed to confirm the robustness of the approach on different cohorts and hospitals.

This work aimed mainly at investigating a new framework for the integration of radiomics and deep learning; limited effort was focused on tuning the DL model, and we restricted the types of radiomics features to those proposed in the reference paper [38]. A similar combination approach of deep and radiomic features has been applied on a subset of the HN dataset to predict distant metastasis by applying CNNs to CT scans only [40]. In particular, we expect that better accuracy can be achieved by adopting specific deep learning architectures or considering more complex methods to extract radiomics features, for instance applying Wavelet filters [17].

## Acknowledgments

The authors thank the WebValley2018 Students Team for initial development of the radiomics environment. The project has been motivated by a discussion on CT/PET modeling with M. Farsad and A. Fracchetti. Part of this work has been supported by the Microsoft Azure Research Award “Deep Learning for Precision Medicine”, assigned to CF.

## Footnotes

1 publicly available at https://wiki.cancerimagingarchive.net/display/Public/ Head-Neck-PET-CT

